# Distribution, Prevalence of Non-Tuberculous Mycobacteria on Hainan Island and Antibiotic Resistance of *Mycobacterium abscessus*

**DOI:** 10.1101/2024.06.09.596927

**Authors:** Xinru Xu, Jingjing Huang, Hongkun Wang, Tianchen Xiang, Jun Liu, Li Zheng

## Abstract

Non-tuberculous mycobacteria (NTM) infections are increasingly recognized as a public health concern, yet their distribution and antibiotic resistance patterns on Hainan Island, China’s tropical hotspot, remain understudied. This study analyzes 158 clinical respiratory samples from 2015–2018 across 14 cities, identifying 118 NTM isolates from 24 species. Predominant species include Mycobacterium abscessus (23.73%), M. intracellulare (22.03%), and M. simiae (14.41%). Spatial mapping reveals elevated prevalence in coastal areas, with no significant north-south divergence on the island (Bray-Curtis dissimilarity: 0.5954, *P* = 0.694), but substantial differences from mainland China (North: 0.9604, *P* = 0.0011; South: 0.9340, *P* = 0.0013). Antimicrobial testing of M. abscessus isolates indicates complete resistance to doxycycline (100%), moderate sensitivity to amikacin (71.4%) and linezolid (50.0%), and variable responses to other agents. Novel detections of M. algericus and M. sinensis expand Hainan’s NTM diversity. These findings establish a baseline for region-specific surveillance, antibiotic stewardship, and intervention strategies to combat NTM misdiagnosis and mortality in tropical settings.

**IMPORTANCE:** Previous studies have not covered the distribution of NTM in various regions of Hainan province. This study provides the first comprehensive geographic map of NTM in Hainan, China’s tropical island hotspot. By monitoring NTM across 14 cities over four years, we reveal key patterns in their habitat preferences and dominant species in infections. These findings establish a foundational framework for region-specific surveillance and treatment protocols in Hainan and offer a replicable model for global tropical regions combating NTM infections. Ultimately, this work empowers public health initiatives to reduce misdiagnosis, enhance antibiotic stewardship, and lower mortality rates among at-risk populations.

## INTRODUCTION

Non-tuberculous mycobacteria (NTM), belonging to the Mycobacteriaceae family, ubiquitously inhabit natural environments such as soil, water, and even the human body. With over 200 documented species, of which approximately 60 are pathogenic, these organisms can infect diverse tissues, including skin, soft tissues, lymphatics, synovial fluid, joints, bones, and lungs (1). In immunocompromised individuals, pathogenic NTM can lead to opportunistic NTM pulmonary disease (NTM-PD) and extrapulmonary infections(2). The global incidence of NTM infections has been increasing, particularly in developing countries (3). In China, the prevalence of NTM infections rose from 4.9% in 1990 to 22.9% in 2010, with the proportion of NTM-related lung infections increasing from 4.24% to 12.68% by 2021 (4, 5). Misdiagnosis of NTM as multidrug-resistant tuberculosis or co-infections with other chronic diseases contributes to escalating mortality rates(6, 7). However, due to the absence of mandatory reporting systems for NTM infections in many nations, current epidemiological data is limited and potentially underestimates the actual burden(8, 9). An accurate understanding of NTM epidemiology and the implementation of effective surveillance systems are thus crucial for guiding control strategies.

Most regions in China have established comprehensive monitoring systems and research strategies for NTM, especially in developed areas like Shanghai, Beijing, and Guangdong, which feature extensive temporal coverage, diverse sample collection, and a wealth of NTM subspecies, leading to well-developed theoretical frameworks and even rare identifications (10–14). NTM research of Hainan Island, initiated in 2019(15), has faced challenges such as geographical limitations and insufficient sample diversity, resulting in a significant lag compared to other provinces. This study aims to enhance the breadth and depth of NTM investigation in Hainan, providing a robust foundation for more precise and targeted intervention strategies, ensuring their scientific rigor and effectiveness, and advancing the scientific frontiers in NTM prevention and control.

*Mycobacterium avium* complex (MAC), the most common slowly growing mycobacteria (SGM), is the predominant cause of NTM pulmonary disease globally(16), with strains like *M. avium, M. intracellulare, M. chimaera, M. marseillense, M. bouchedurhonense, M. timonense, M. yongonense*, and *M. paraintracellulare* recognized as pathogens (17). The *M. abscessus* complex (MABC), dominated by *M. abscessus* subspecies, accounts for 65%-80% of rapidly growing mycobacteria (RGM) pulmonary infections(18) (19), with three key subspecies identified. *M. kansasii* ranks as the second most frequent NTM pathogen in some European countries(20), whereas in China, MAC and MABC prevail, specifically *M. intracellulare* and *M. abscessus*, along with other pathogens such as *M. fortuitum, M. gordonae*, and *M. kansasii*. In China, NTM distributions exhibit a southern, coastal, and warm-humid regional predominance over northern, inland, and cold-dry areas(2, 5).

Drug susceptibility testing of numerous NTM isolates indicates clarithromycin (CLR) and amikacin (AMK) as highly effective against MAC, serving as first-line agents. MAC exhibits resistance to linezolid (42.86%) and moxifloxacin (25.32%), positioning these as second-line options.. Avoiding drugs like those initially can prevent ineffective treatment.

MABC exhibits high resistance to moxifloxacin, ciprofloxacin, and tobramycin (>95%), alerting clinicians not to rely on these drugs for treating MABC-related diseases. Instead, the relatively high susceptibility of MABC to tigecycline (98.53%) and amikacin (64.71%), highlights their potential as first-line therapies(21). This knowledge can prevent the use of ineffective antibiotics, which may waste time and resources and worsen the patient’s condition. Given the suboptimal outcomes of current NTM treatments, the exploration of alternative antibiotics such as clofazimine, bedaquiline, delamanid, and tedizolid is warranted. These drugs have shown in vitro activity against specific NTM species, including *M. intracellulare, M. avium* (clofazimine-sensitive), *M. abscessus, M. fortuitum*, and *M. chelonae* (tedizolid-responsive), highlighting their therapeutic potential (22).

Prevots’ research(23) identified atmospheric hydrologic parameters as critical drivers of non-tuberculous mycobacteria (NTM) ecological transmission dynamics, with significant spatial associations between aerosol-phase water loading and pathogen prevalence. In Hainan Island, the observed biogeographic divergence of NTM species clusters across climate gradients underscores the need for region-adapted frameworks to address possible tropical maritime environmental determinants. Previous studies have not covered the distribution of NTM in various regions of Hainan province. We delve into the distribution and antibiotic resistance profiles of NTM in Hainan, elucidating the current NTM prevalence scenario and augmenting the province’s NTM research corpus.

## METHODS

### Ethics Statement

This study was conducted with the approval of the Ethics Committee of Hainan Medical University, under the ethics number HYLL-2021-170.

### Study Population and Sample Collection

The experimental samples were sourced from the Hainan Tuberculosis Prevention and Control Institution, covering the period 2015-2018. A total of 158 sputum and bronchoalveolar lavage fluid specimens, clinically suspected of NTM infection, were collected from 14 cities and counties in Hainan Island, including Baisha, Baoting, Changjiang, Chengmai, Danzhou, Ding’an, Dongfang, Haikou, Ledong, Lingao, Lingshui, Qionghai, Sanya, and Wenchang.

### Materials and Equipment

Bacterial genomic DNA extraction kit (Tiangen Biotech, Beijing), Template DNA, buffer, dNTPs, TaqDNA polymerase (Tiangen Biotech, Beijing), Hsp65 and rpoB gene amplification primers (Tsingke Biotech, Beijing), PCR thermal cycler (Bio-Rad Laboratories, Shanghai), DNA recovery kit (Tiangen Biotech, Beijing). High-speed benchtop centrifuge (Eppendorf, Germany).

### Clinical Sample Collection

Patients were instructed to fast for at least 1-2 hours before sample collection. Before expectoration, patients perform a mouthwash with 0.1% spirulina solution three times, followed by rinsing with clean water, to minimize oral flora contamination. Sputum Production: Deep coughing is encouraged to ensure productive sputum collection into sterile plastic containers with a screw-mouth, provided by the hospital, emphasizing the importance of collecting deep sputum such as purulent, mucoid, or blood-streaked specimens. Saliva samples may also be accepted if no productive sputum can be produced. Each patient provides three types of sputum samples (morning, night, and on-site) over a day. Morning and night sputa are self-collected at home, while the on-site sample is obtained at the hospital. Samples are transported to the tuberculosis laboratory within 4 hours of collection. If immediate transportation is not feasible, samples are refrigerated at 4°C until delivery. In the absence of a provincial tuberculosis control institute, samples are sent to designated tuberculosis laboratories in each city for initial testing. Subsequently, positive cultures are forwarded to the Hainan Provincial Center for Disease Control and Prevention for confirmation and further analysis.

### PCR-based Species Identification

DNA was extracted from each sample to serve as a template for subsequent PCR reactions using primers targeting rpoB and Hsp65 genes. Primer sequences were as follows: Hsp65-F: 5’-ACC AAC GAT GGT GTG TCC AT-3’, Hsp65-R: 5’-CTT GTC GAA CCG CAT ACC CT-3’, rpoB-F: 5’-GAC GAC ATC GAC CAC TTC GG -3’, rpoB-R: 5’-GGG GTC TCG ATC GGG CAC AT-3’. Amplification was conducted in 20 μL reaction mixtures containing 2 μL 10×PCR buffer, 200 μM dNTPs, 0.25 μM of each primer, and 1U TransTaq HiFi DNA polymerase (Tiangen Biotech, Beijing). Thermal cycling conditions comprised initial denaturation at 94°C for 5 minutes; 30 cycles of 94°C denaturation for 30 seconds, annealing at 60°C for 30 seconds, extension at 72°C for 1 minute; and a final extension at 72°C for 10 minutes. Products were analyzed by agarose gel electrophoresis, and DNA fragments were recovered using a gel extraction kit and sent to Hainan Nanshan Biotechnology Co., Ltd. for sequencing. Ultimately, nucleotide sequences were aligned with GenBank references via the NCBI BLAST website on 04/22/2024 (https://blast.ncbi.nlm.nih.gov/Blast.cg).

### Phylogenetic Tree Construction

To explore evolutionary relationships between Hsp65 and rpoB genes, multiple sequence alignment was performed using the Muscle algorithm in MEGA software (v11.0) with default parameters. Neighbor-joining (NJ) method was applied in MEGA to construct phylogenetic trees. To enhance tree topology reliability, a bootstrapping process involving 1000 iterations was executed. For intuitive comprehension, the phylogenetic tree was refined and annotated using the Tree Visualization by One Table (TvBOT, https://www.chiplot.online/tvbot.html, version 2.6) online tool.

### Drug Sensitivity Test

According to Clinical and Laboratory Standards Institute (CLSI) guidelines, broth microdilution assays were employed for in vitro antimicrobial susceptibility testing of selected NTM isolates. Minimum inhibitory concentration (MIC) values require clinical context interpretation and adjustment. MIC50 is defined as the lowest concentration of the drug that inhibits the growth of 50% of the tested M. abscessus isolates; MIC90 is the lowest concentration that inhibits 90% of the tested *M. abscessus* isolates. The MIC breakpoints used to define susceptibility categories are based on the Clinical and Laboratory Standards Institute (CLSI) guidelines, specifically the M24-A2 document, with the exception of azithromycin. For amikacin, linezolid, moxifloxacin, and doxycycline, the breakpoints are as follows: Amikacin: Susceptible (S) ≤ 16 μg/ml, Intermediate (I) 32 μg/ml, Resistant (R) ≥ 64 μg/ml. Linezolid: S ≤ 8 μg/ml, I 16 μg/ml, R ≥ 32 μg/ml; Moxifloxacin: S ≤ 1 μg/ml, I 2 μg/ml, R ≥ 4 μg/ml; Doxycycline: S ≤ 1 μg/ml, I 2 - 4 μg/ml, R ≥ 8 μg/ml. These breakpoints are based on CLSI M24 - A2(24). For azithromycin, the breakpoints (S ≤ 2 μg/ml, I 4 μg/ml, R ≥ 8 μg/ml) are derived from Li,2017(25), as there is no direct CLSI breakpoint for this drug against M. abscessus.

The MicroDST™ system (Yinked AUTOBIO Diagnostics Co., Ltd., Zhuhai, China) was utilized for resistance testing of NTM isolates. MIC breakpoints and sensitivity/resistance determination were interpreted per manufacturer instructions. Five antibiotics were tested against *M. abscessus* isolates: moxifloxacin, amikacin, linezolid, azithromycin, and doxycycline.

### Data Analysis

Bray-Curtis dissimilarity matrices were computed via the vegdist function (vegan package in R) after data preprocessing (removing all-zero species/samples, transposing matrices). Permutational Multivariate Analysis of Variance (PERMANOVA) was conducted to test for significant differences in NTM species composition between regions using adonis2 with 9999 permutations. A non-parametric bootstrap (10000 replications) with the bias-corrected and accelerated (BCa) method estimated the 95% confidence interval for the mean inter-group Bray-Curtis distance.

## RESULTS

### NTM Subspecies Characterization

From 158 clinical samples suspected of NTM collected between 2015 and 2018,157 isolates were successfully species-identified, comprising 118 NTM (across 24 species) and 39 non-NTM organisms. Identification primarily relied on rpoB gene sequencing, with Hsp65 sequencing serving as a supplementary tool for conclusive determination. All data have been uploaded to https://nmdc.cn/submit/dashboard: Hsp65 (NMDCN0007ON9 to NMDCN0007OQQ) and rpoB (NMDCN0007OQR to NMDCN0007OUG). The species distribution, as shown in Figure 1, was led by *M. abscessus* (28/118, 23.73%), *M. intracellulare* (26/118, 22.03%), *M. simiae* (17/118, 14.41%), *M. paraintracellulare* (16/118, 13.56%), and *M. colombiense* (5/118, 4.24%). MTB(23), *Nocardia thailandica* (3/16, 18.75%), *Tsukamurella paurometabola* (2/16, 12.5%), and *Gordonia jinghuaiqii* (2/16, 12.5%), were also identified. Detailed composition ratios are illustrated in Table 1.

**Table 1.**
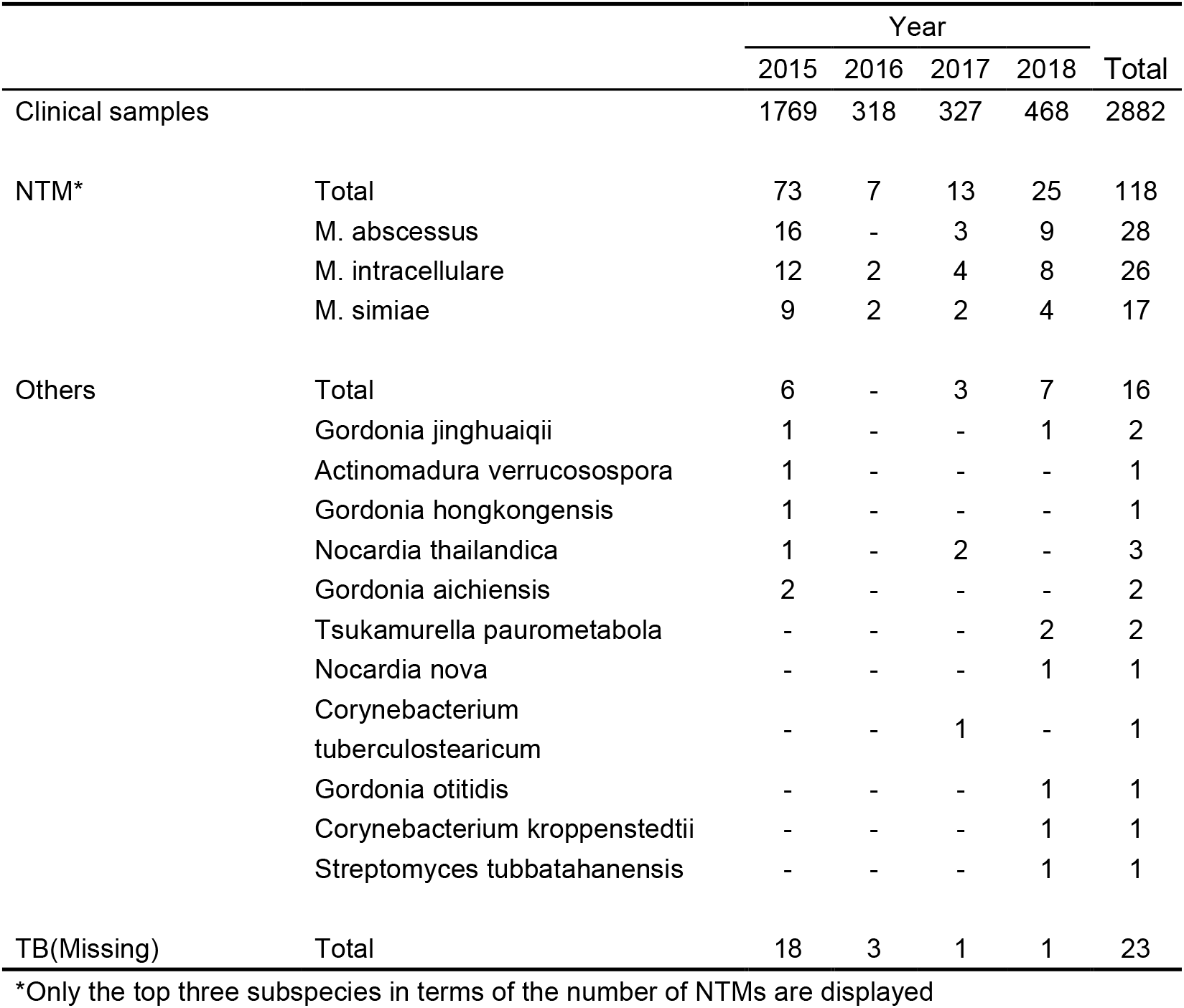
NTM species identified from 2015 to 2018 in Hainan, China.

**Figure 1.**
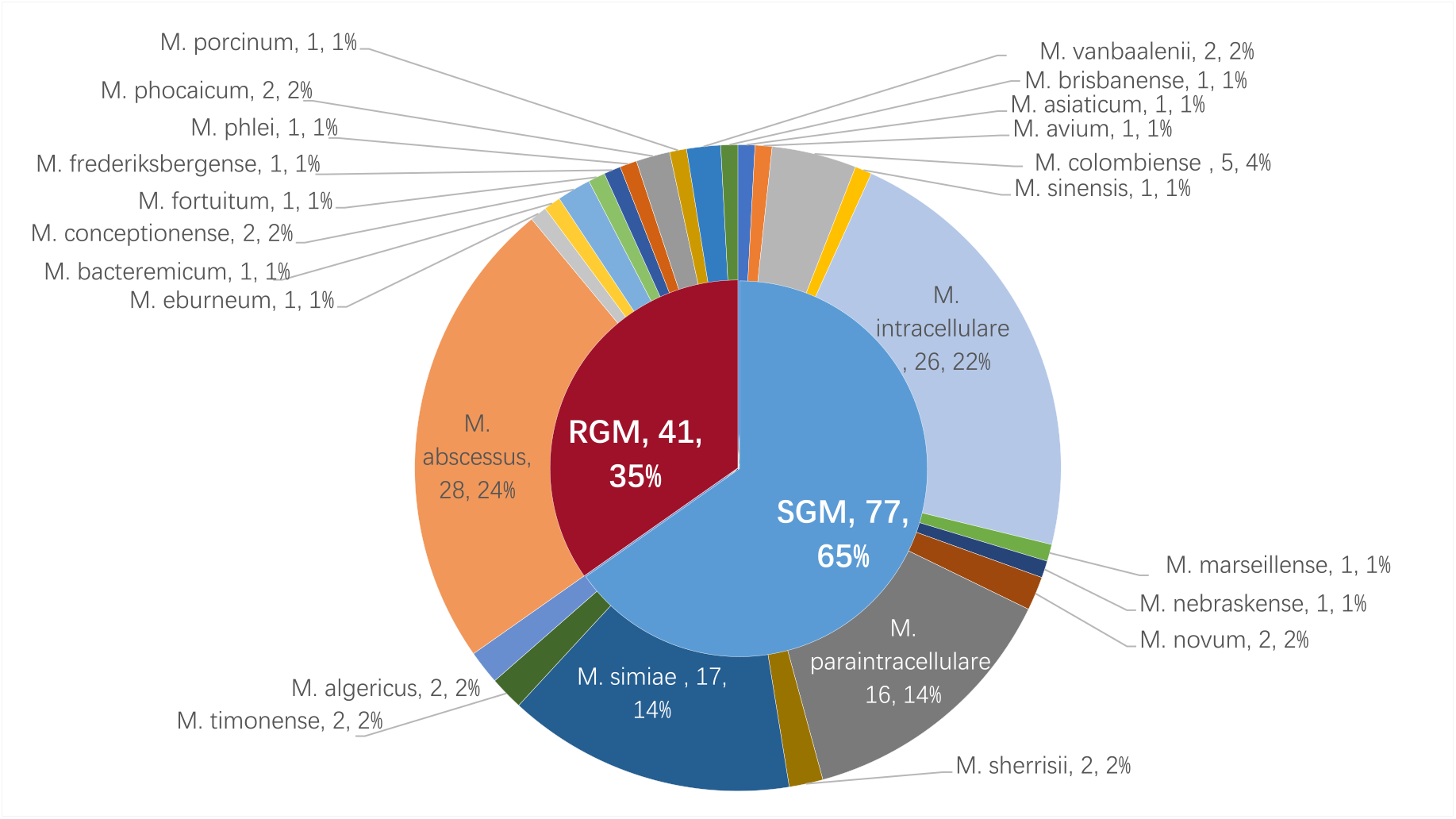
Distribution of non-tuberculous Mycobacteria isolated from patients with suspected pulmonary NTM in Hainan Island, China. Abbreviation: NTM: non-tuberculous Mycobacteria. RGM: rapidly growing mycobacteria; SGM: slowly growing mycobacteria.

### rpoB Gene Phylogeny of NTM

A neighbor-joining (NJ) phylogenetic tree, based on NTM rpoB sequences, revealed distinct clustering patterns among pathogenic and potentially pathogenic species. *M. fortuitum* and *M. abscessus* grouped into a distinct cluster of rapidly growing, pathogenic mycobacteria. Strong bootstrap support validated the tree’s reliability, with *M. sherrisii* and *M. simiae* closely related per Hsp65 and rpoB analysis, reflecting proximity to MAC, MABC, and *M. simiae* complexes(Figure 2). Hsp65 Gene Phylogeny of NTM was shown in Supplemental Figure 1.

**Figure 2.**
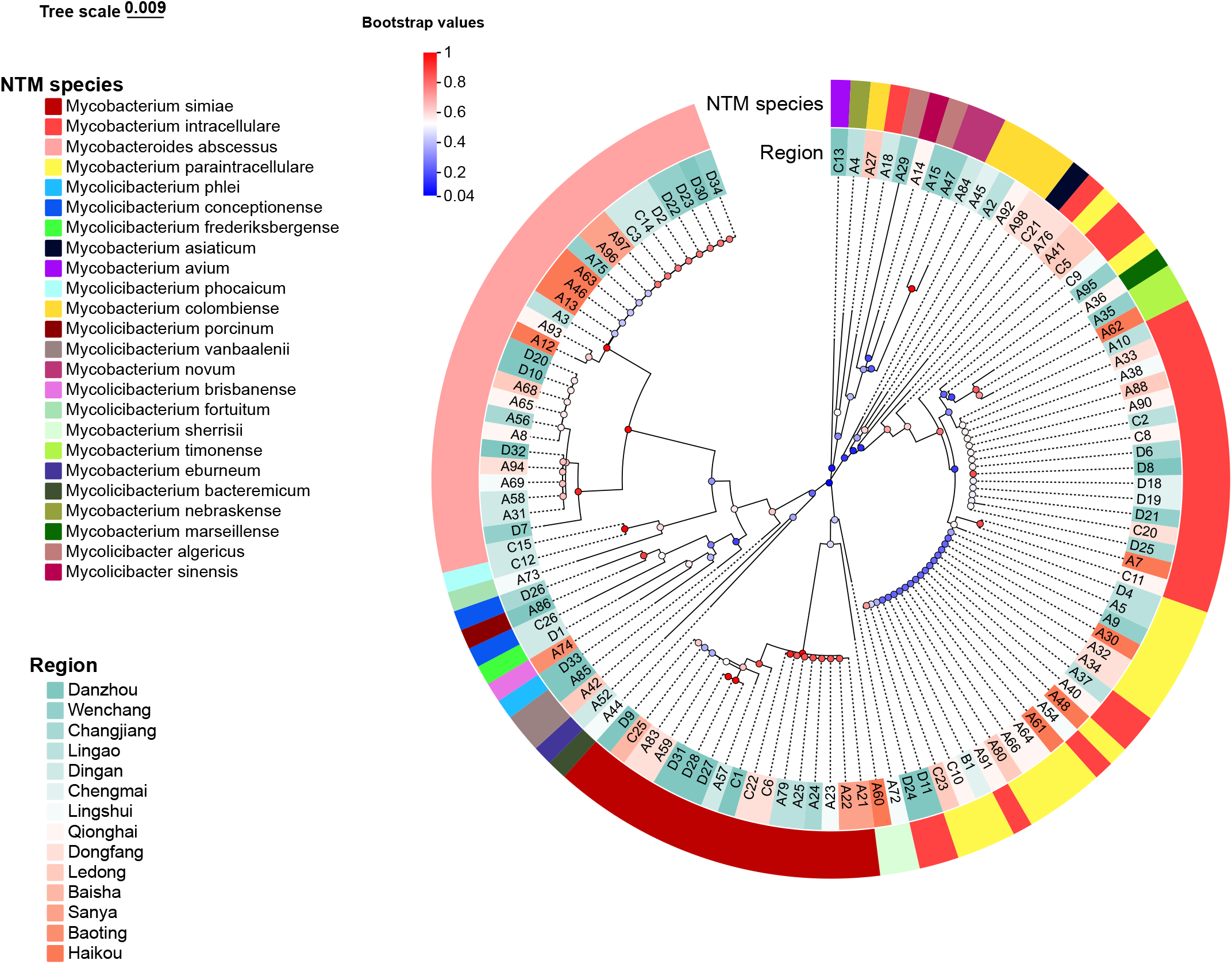
Phylogenetic tree of rpoB gene from 118 NTM using the neighbor-joining method with MEGA11.0 software. Tree-scale 0.007 represents the evolutionary distance unit, and Bootstrap values represents the self-expanding value, which is used to test the calculated evolutionary Tree branch confidence.

### Spatial Distribution of NTM in Hainan Island

The 118 NTM isolates were distributed across 14 cities in Hainan, with distinct patterns. Northeastern regions reported higher NTM presence compared to southwestern, and SGMs outnumbered RGMs. MAC and MABC were dominant NTM groups, featuring *M. intracellulare* and *M. abscessus* respectively. SGMs were detected in all except Baoting, while RGMs were absent in Chengmai and Baisha. Key distributions included *M. intracellulare* in Qionghai, Lingshui, Dongfang, and Danzhou, and *M. abscessus* in Dingan, Wenchang, and Haikou. Species like *M. algericus, M. sinensis, M. bacteremicum*, and others showed unique distributions. Refer to Figure 3 for comprehensive mapping.

**Figure 3.**
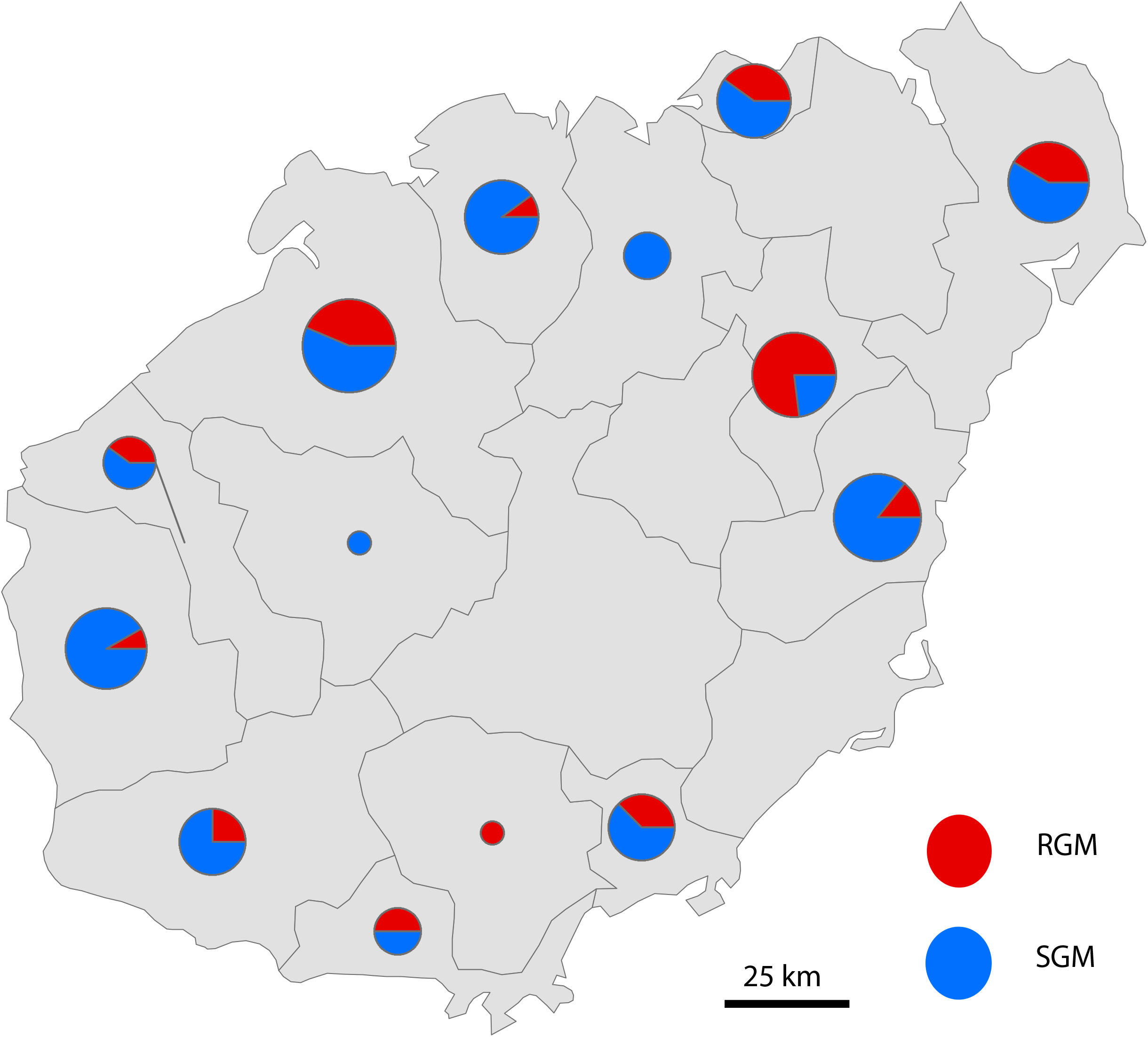
Distribution map of NTM in 14 cities and counties in Hainan Province (Baisha, Baoting, Changjiang, Chengmai, Danzhou, Ding’an, Dongfang, Haikou, Ledong, Lingao, Lingshui, Qionghai, Sanya, Wenchang).

### Regional Comparison

Bray-Curtis dissimilarity analysis revealed three distinct epidemiological clusters (Table 2). Intra-island variation: The northern and southern regions of Hainan Island demonstrated moderate microbial differentiation (BC = 0.5954, *P* = 0.694), which indicates that the difference in the NTM strain communities between the northern and southern parts within Hainan Province is not significant (Details in TABLE S1). Inter-regional divergence: Striking contrasts emerged between Hainan Island and mainland China, with the dissimilarity between Hainan and North China reaching 0.9604 (P =0.0011). Hainan-South China dissimilarity measured 0.9340 (*P* = 0.0013) (Details in TABLE S2).

**Table 2.** Bray-Curtis Dissimilarity Analysis of NTM Strain.

| Region Comparison | Bray-Curtis Distance | *P* | 95%CI |
| --- | --- | --- | --- |
| Northern Hainan vs. Southern Hainan | 0.5954 | 0.694 | [0.5234, 0.6672] |
| North (CHINA)(26–28)vs. Hainan | 0.9604 | 0.0011 | [0.9491, 0.9688] |
| South (CHINA)(29–31) vs. Hainan | 0.9340 | 0.0013 | [0.9124, 0.9499] |

### Susceptibility Test Results of *M. abscessus*

The MICs and breakpoints for seven antibiotics against 28 *M. abscessus* isolates are detailed in Table 3. The isolates displayed high levels of antibiotic resistance, with universal resistance to doxycycline but retained sensitivity to amikacin (71.4%) and linezolid (50.0%). Complete resistance and sensitivity profiles are elaborated in Table 3.

**Table 3.** Drug susceptibility test of *M*.*abscessus* (n=28)

| Antimicrobial agent | MIC(μg/mL) |  |  |  |  |  | Number(%) |  |  |
| --- | --- | --- | --- | --- | --- | --- | --- | --- | --- |
|  | MIC range | MIC 50 | MIC 90 | S | I | R | S | I | R |
| Amikacin | <1-64 | 4 | 64 | ≤16 | 32 | ≥64 | (20/28)71.4% | (3/28)10.7% | (5/28)17.9% |
| Linezolid | 2-32 | 8 | 32 | ≤8 | 16 | ≥32 | (14/28)50.0% | (8/28)28.6% | (6/28)21.4% |
| Moxifloxacin | 0.125-16 | 4 | 16 | ≤1 | 2 | ≥4 | (8/28)28.6% | (6/28)21.4% | (14/28)50.0% |
| Azithromycin | <1-32 | 16 | 32 | ≤2 | 4 | ≥8 | (9/28)32.1% | (5/28)17.9% | (14/28)50.0% |
| Doxycycline | 16->128 | >128 | >128 | ≤1 | 2~4 | ≥8 | 0 | 0 | (28/28)100% |

## DISCUSSION

This study identified five major groups of NTM: MAC, MABC, *M. simiae* complex, *M. fortuitum* complex, and other NTM species. MAC, a common causative agent of human NTM diseases affecting various organs such as lungs, soft tissues, lymph nodes, and bone marrow, with MAC pulmonary disease being the most prevalent(21), was also the primary NTM pathogen detected in our study from pulmonary disease cases in Hainan, with a total of 51 isolates (comprising *M. intracellulare, M. paraintracellulare, M. colombiense, M. timonense, M. avium*). This aligns with previous reports by Zhong (15) and Wang (21) who identified 149 and 154 MAC isolates respectively, reinforcing the widespread distribution of MAC in Hainan. MABC, another dominant NTM group, predominantly consisted of *M. abscessus* and its subspecies, differing slightly from Zhong and Wang’s findings where *M. intracellulare* was more frequency. *M. simiae* complex emerged as the third most frequent NTM in Hainan, further supporting its presence as a significant species in the region, contrasting its less frequent detection in other provinces(2). The NTM profile in Hainan generally aligns with previous studies by Zhong (15, 32),Wang(21),and Wu(33), showing a higher prevalence of SGMs compared to RGMs. Huang(34) in Beijing also found *M. intracellulare* as the dominant NTM, followed by *M. abscessus* and *M. avium*, mirroring our results where *M. abscessus* and *M. intracellulare* dominate.

No statistically significant difference in NTM strain composition was observed between northern and southern Hainan Island. This similarity is driven by two key factors: both regions share a tropical monsoon climate, lacking sufficient climatic variability to drive NTM species differentiation; human activity patterns are highly uniform, with tourism and tropical agriculture dominating economic activities; and no significant social drivers exist to influence NTM transmission dynamics. However, this study identified Danzhou as the region with the highest NTM isolation frequency, contrasting with earlier studies (2015–2018) that reported Haikou as the primary hotspot(15). This discrepancy arises from methodological advancements, in which our use of Hsp65 and rpoB gene sequencing, combined with GenBank sequence alignment, enabled precise identification of NTM subspecies, Mycobacterium tuberculosis (MTB), and contaminating bacteria, unlike prior studies that relied on gene chip technology limited to detecting 17 common mycobacterial species. These improvements enhance the thoroughness and reliability of our findings.

A marked difference in NTM strain composition was observed between northern mainland China(26–28) and Hainan Island (BC = 0.9604), reflecting distinct regional NTM profiles. In Hainan, M. abscessus predominates, favored by the region’s hot, humid tropical climate, which is less suitable for M. intracellulare, the dominant NTM in northern China’s drier environments, such as Beijing(26), where it accounts for 40–60% of isolates(35). Northern regions typically exhibit a higher prevalence of slow-growing mycobacteria (SGM), with M. intracellulare as a key representative, compared to rapidly growing mycobacteria (RGM) like M. abscessus. Additionally, regional disparities extend to less common species: Mycobacterium lentiflavum, the second most frequently isolated NTM in Tianjin, was absent in Hainan, underscoring the geographic specificity of NTM distributions across China.

Significant differences in NTM strain composition were also found between southern mainland China(29–31) and Hainan Island (BC = 0.9340). In southern China(36), *M. abscessus* is predominance, consistent with its dominance in Hainan, where it was the most frequently isolated NTM. However, species such as *M. kansasii* and *Mycobacterium smegmatis*, detected elsewhere in China, were absent in Hainan. In contrast, *M. avium* predominates in regions like Sichuan(37) and coastal southeastern provinces, e.g., Fujian(38), Zhejiang(39), Jiangsu(40), where it is more frequently isolated than in Hainan, which reported only one single M. avium isolate. These findings highlight Hainan’s unique NTM assemblage, shaped by its distinct environmental and geographic characteristics, distinguishing it from both northern and southern mainland China.

Considering Hainan and Taiwan’s insular geography and similar latitudes, comparing NTM distributions reveals insights. Huang et al. (41) found *M. abscessus* and MAC to be the most common in Taiwan’s NTM-pulmonary infections, mirroring Hainan’s MAC dominance and *M. abscessus* prevalence. Although no significant statistical difference was found, subtle variations may exist and warrant further investigation.Hainan’s northern NTM abundance contrasts with Taiwan’s southern predominance, possibly influenced by Hainan’s latitude, tropical/subtropical climates, and topography. Hainan’s central mountainous terrain might influence coastal NTM prevalence, reflecting the island’s central-high and peripheral-low geography. Overall, these findings underscore the impact of geographical and environmental factors on NTM distribution patterns across regions.

As opportunistic pathogens, NTM typically invade organs such as the lungs, skin, lymph nodes, and bones when host immunity is compromised, with lung infections being most common(42, 43). Although NTM exhibits lower virulence and pathogenicity than MTB, their non-specific clinical symptoms and radiological similarities to MTB lead to diagnostic challenges. Their natural and acquired drug resistance further complicates clinical management(43) underscoring the importance of NTM species identification and epidemiological investigations.

In Hainan, the hot and humid environment fosters NTM survival and dissemination, making epidemiological studies in this high NTM-endemic region crucial for local control efforts. Given *M. abscessus*’s natural and acquired resistance to many common antibiotics, resistance testing to identify effective treatments is essential; clinical practice often employs anti-tubercular drugs(44). Guidelines(45) recommend clarithromycin, azithromycin, amikacin, and cefoxitin for *M. abscessus*. Our susceptibility tests revealed rates of 71.4% for amikacin, 50.0% for linezolid, 32.1% for azithromycin, and 28.6% for moxifloxacin. The high resistance to doxycycline (100%) highlights the need for caution when using this drug clinically. The intrinsic resistance mechanisms of *M. abscessus* include its waxy, opaque cell wall, drug transport systems, antibiotic modification, and target gene polymorphisms(46). Subspecies differences in resistance rates (47), particularly with higher clarithromycin resistance in *M. abscessus subsp. abscessus* than *subsp. massiliense*, indicating a need for further study to guide treatment.

Comparative phylogenetic analysis reveals distinct evolutionary trajectories between Hsp65 and rpoB genes in nontuberculous mycobacteria (NTM): The Hsp65 tree exhibits uniform topology with conserved sequence architecture ideal for resolving macroevolutionary relationships(48), whereas the rpoB phylogeny displays accelerated evolutionary rates and intricate branching patterns that facilitate high-resolution discrimination of closely related strains(49). Hsp65-based phylogenetic analysis demonstrated limited resolution in differentiating *Mycobacterium intracellulare* from *M. paraintracellulare*, resulting in misidentification of 16/49 (32.7%) *M. paraintracellulare* strains as *M. intracellulare*. In contrast, rpoB gene sequencing achieved unambiguous discrimination between these two species. Both phylogenetic trees revealed close evolutionary proximity between *M. sherrisii* and *M. simiae*, though with divergent statistical support: strong bootstrap values (0.95) in the Hsp65 tree versus moderate support (0.45) in the rpoB phylogeny, indicating Hsp65’s superior discriminative capacity for this species pair. Notably, weak bootstrap support (for the *M. sherrisii*–*M. simiae* clade in the rpoB tree) does not invalidate their inferred evolutionary relationship, as this conclusion is corroborated by two complementary lines of evidence: (1) strong nodal support (0.95) for the same clade in the Hsp65 phylogeny;(2) ≥99% rpoB sequence identity between our isolates and the GenBank-retrieved type strains of *M. sherrisii* and *M. simiae*. This necessitates critical awareness during diagnostic interpretation to prevent taxonomic misclassification(50). The rpoB phylogeny further resolved evolutionary relationships within the MAC(51), demonstrating shared ancestral origins (bootstrap: 0.4-1.0) among *M. intracellulare, M. paraintracellulare, M. timonense, M. colombiense*, and *M. marseillense*. This phylogenetic evidence corroborates rpoB’s diagnostic reliability for MAC species delineation, aligning with established multilocus sequence analysis benchmarks. Phylogenetic discordance is observed in rpoB-based clustering of four Mycobacteroides abscessus isolates from Danzhou, which segregate into two distinct clades with robust nodal support (Clade A: 1 isolate, bootstrap=0.98; Clade B: 3 isolates, bootstrap=0.93), despite exhibiting identical mean pairwise distance (0.01 substitutions/site). This polyphyletic clustering suggests ecological selection pressures have driven substantial genomic divergence while maintaining taxonomic coherence under current species delineation criteria.

In recent years, the increasingly widespread whole-genome sequencing (WGS) — a high-precision tool for identifying and characterizing mycobacteria — outperforms techniques like targeted PCR genotyping with superior resolution, enabling differentiation of NTM subspecies into specific clades, providing comprehensive genomic data, and offering unparalleled accuracy in detecting genetic variations, identifying new species, and predicting virulence genes, thus underpinning targeted therapy(52, 53).

The study unexpectedly identified numerous non-NTM species, including previously undetected MTB and species of Corynebacterium, Actinomadura, Gordonia, Tsukamurella, Streptomyces, and Nocardia, all belonging to the actinobacteria phylum. Streptomyces tubbatahanensis, not previously reported in China, suggests new perspectives in streptomycete pathogenesis. Tsukamurella paurometabola, rarely pathogenic but previously isolated in Hainan (21) indicates its opportunistic nature. Nocardia species, ubiquitous yet some highly pathogenic especially in immunocompromised individuals, with Nocardia nova known as the second most prevalent (54), and is known to cause pulmonary infections, while *Nocardia thailandica* has also been documented to induce pneumonia(55). Gordonia species, isolated from various environments, can cause opportunistic infections in immunocompromised hosts. Corynebacterium tuberculostearicum is omnipresent on human skin, yet its health impacts require further investigation(56). Definitive differentiation between contamination and bona fide infection necessitates additional iterative sampling and molecular typing, thereby highlighting the imperative of expanding pathogen surveillance in complex clinical scenarios.

Between 2015 and 2018, the present study documented that Mycobacterium abscessus persisted as the dominant species, without notable alterations. Following the nationwide rollout of Xpert MTB/RIF® (Cepheid) nucleic acid amplification testing in 2016, clinical laboratories systematically discontinued routine submission of acid-fast bacilli (AFB) smear-negative specimens to tuberculosis lab for confirmatory culture. Consequently, the sample size decreased. 2015 data reflects pre-intervention sampling, while subsequent years show the impact of policy changes.

As an expansion and deepening of previous work in Hainan, this study leverages the province-wide TB control network for multi-dimensional NTM testing, achieving notable progress in geographic coverage and species diversity. By involving hospitals at all levels and standardizing specimen submission, a comprehensive and representative NTM repository was established, surpassing single-center biases and enhancing sample representativeness. This collaborative, cross-regional approach lays a firm foundation for elucidating regional NTM patterns and potential epidemiological rules. It also underscores the direct influence of public health policies on monitoring activities, emphasizing the need for flexibility and adaptability in future surveillance systems to accurately track NTM epidemiology. Collectively, this study enriches Hainan’s NTM epidemiological data, informing targeted control strategies, and highlighting the value of interdisciplinary and inter-institutional cooperation in addressing NTM public health challenges.

## CONCLUSION

This study has provided an in-depth exploration into the geographic distribution and drug resistance patterns of NTM in Hainan Island, highlighting *M. abscessus* and MAC as principal pathogens driving NTM infections in the region, predominantly in Danzhou, Qionghai, Dingan. Novel findings include the documentation of previously unreported NTM species in Hainan, specifically *M. novum*; *M. algericus*; *M. conceptionense*; *M. vanbaalenii*; *M. eburneum*; *M. nebraskense*; *M. sinensis*; *M. bacteremicum*; *M. frederiksbergense*; *M. phlei*. Moreover, the study underscores the therapeutic potential of amikacin and linezolid against *M. abscessus*.

## DATA AVAILABILITY

All sequencing data generated in this study have been deposited in the National Microbiology Data Center of China. The sequencing results of Hsp65 genes have been assigned accession numbers ranging from NMDCN0007ON9 to NMDCN0007OQQ, and rpoB genes from NMDCN0007OQR to NMDCN0007OUG.

## AUTHOR CONTRIBUTION

XX, HW, and TX conducted the experiments; JH was responsible for sample collection; XX, JL, and JH performed the data analysis; XX and LZ wrote and revised the manuscript; LZ designed the study.

